# Hi-C sequencing data from cortex of laboratory rats

**DOI:** 10.1101/2025.02.09.636698

**Authors:** Panjun Kim, Rachel R. Ward, Burt M. Sharp, Robert W. Williams, Hao Chen

**Author notes:** Correspondence: Hao Chen, 71. S. Manassas Street, Room 209 Translational Science Building, University of Tennessee Health Science Center, Memphis, TN 38103.

## Abstract

The three-dimensional conformation and packaging of chromosomes modulates the spatial organization of the nucleus, orchestrating DNA replication and repair, maintaining genome stability and integrity, and regulating gene expression. Hi-C methods provide high-resolution data on chromatin-to-chromatin interactions both within and among chromosomes at a genome-wide scale. Hi-C resolves topologically-associated domains (TADs) and chromatin loops that are linked to cell-specific transcriptional control. We present a comprehensive Hi-C dataset generated from the frontal cortex of laboratory rats, encompassing a diverse group of inbred strains (SHR/OlaIpcv, BN-Lx/Cub, BXH6/Cub, HXB2/Ipcv, HXB10/Ipcv, HXB23/Ipcv, HXB31/Ipcv, LE/Stm, F344/Stm) and an F1 hybrid (SHR/Olalpcv x BN/NHsdMcwi). This dataset serves as a valuable resource for studying the mechanisms by which three-dimensional chromatin architecture governs gene expression in the brain. Strain-specific variations in genome organization can illuminate the influence of chromatin structure on gene expression, neuronal functionality, and the predisposition to neurological and behavioral disorders.

## Background & Summary

To understand 3D genome architecture within the nucleus, imaging and Chromosome Conformation Capture (3C) led the research for 3D genome structure^1^. Hi-C sequencing, derived from the combination of the 3C assay combined with high-throughput Next-Generation Sequencing (NGS) technology, has become widely utilized due to their ability to map chromatin interactions across the genome at a high resolution^2^. The chromatin contact frequencies obtained through Hi-C revealed the hierarchical 3D structures of the genome, including chromosomal compartments, topologically associated domains (TADs), and chromatin loops. Notably, such structures have been recognized for their significant roles in various biological and cellular processes, including the regulation of gene expression^3^, modulation of chromatin accessibility^3^, and their influence on disease mechanisms^4^. Also, these data on long-range interactions between chromatin segments are valuable for a range of applications in genomics and biomedical research.

A major application of the HI-C method is in genome assembly, where Hi-C data are used to scaffold contigs into chromosome-scale assemblies, enabling the construction of highly contiguous and accurate genome references. This is possible because Hi-C contact frequencies provide spatial proximity data, indicating which genomic fragments are likely to be physically adjacent. By analyzing the interaction data, researchers can accurately order and orient contigs, resolving ambiguities in assembly derived from linear sequencing data. This approach enhances the completeness and accuracy of genome assemblies. This application is particularly beneficial for resolving complex genomes, including those of polyploid species^5^, and for haplotype phasing, which separates homologous chromosomes into distinct assemblies^6^. Additionally, Hi-C data can be exploited to identify structural variants (SVs) providing valuable insights into genomic rearrangements, including deletions, duplications, insertions, inversions, and translocations. These SVs typically involve alterations spanning 50 base pairs or more^7^, making them challenging to detect using short-read sequencing technologies due to their limited capacity for their read length and repetitive region^8^. SVs, such as translocations or inversions, can significantly alter the spatial distance between two genomic regions. These SVs change the contact frequency of reads mapped to these regions. Compared to the background interaction frequencies of individuals without these variants, these altered interactions serve as indicators of SVs^9^.

Chromatin loops identified by Hi-C, in general, are formed by enhancer-promoter interactions, which are essential for understanding transcriptional regulation^10^. In addition, information on promoter-enhancer interactions and structural variations obtained from Hi-C data have become an integral part of multi-omics integration, enabling researchers to better understand regulatory mechanisms in transcriptomic, epigenomic, and proteomic data^11^. This integrative approach provides a comprehensive view of how genome organization influences cellular processes, uncovering relationships between chromatin architecture and functional genomic elements in both health and disease contexts.

## Methods

### Tissue processing, Hi-C library preparation, and sequencing

Frozen rat brains were removed from long-term storage at -80°C, stored at -20°C for ≥ 30 minutes, and cryosectioned at -15 °C into 120 µm slices. Using a -10°C cold block, 30-200 mg of cortex tissue was microdissected using an 18-gauge needle with a bent tip, and a pair of forceps as needed. Optimal tissue input weight was found to be ∼50 mg after extensive troubleshooting. Bilateral cortex tissue was immediately transferred to pre-chilled 1.5 mL DNA Lobind tubes (Eppendorf) kept on dry ice until all tissue was collected. Tissue was pulverized in small quantities using a Cellcrusher mini (Cellcrusher) pre-cooled in liquid nitrogen until all tissue had been pulverized and collected. 1X PBS was used to collect samples from the Cellcrusher. Tissue samples were then processed according to the Arima Hi-C+ Kit (Arima). Samples that passed QC1 in the Arima Hi-C protocol continued to the Arima-provided KAPA Hyper Prep Kit (Roche) protocol. Samples were fragmented using a Covaris S2 instrument (Table 1), generally with the following settings: Duty Cycle 10%, Intensity 5, Cycles/Burst 200, Time 55 s, # cycles 8-15, using Covaris microTUBEs (Covaris). Fragmented samples were analyzed by Agilent Bioanalyzer to confirm a median fragment size of 200-600. Size selection methods varied throughout the samples due to extensive troubleshooting; optimal size selection was performed with fresh Ampure XP beads (Beckman Coulter) in a 2-sided manner, with the goal of a narrow peak and 400 bp median fragment. Any steps that involve Ampure XP beads were performed with either a DynaMag (Thermo Fisher) compatible with 1.5 mL tubes, or a magnetic rack with high/low settings (Parse Biosciences) compatible with PCR-sized tubes; magnetic 96-well plates compatible with PCR-sized tubes were found to result in bead loss. Library preparation was performed using the KAPA HyperPrep kit and Illumina TruSeq DNA Single Indexes (Illumina; discontinued) or Illumina TruSeq UDI v2 (Illumina; current version). Pre-amplification libraries were quantified using the KAPA Library Quantification Kit (Roche) and Lightcycler 480. Amplified libraries were analyzed using Qubit HS DNA Assay (Invitrogen) and Agilent Bioanalyzer HS DNA chip. All other quantification steps in the Arima and KAPA protocols were also performed with the Qubit HS DNA Assay and Agilent Bioanalyzer HS DNA chips. Available part numbers for all listed reagents have been provided in **Table1**.

**Table 1.** Kits and reagents for library preparation.

| Kit/Reagent | Supplier | Part No. | Notes |
| --- | --- | --- | --- |
| Arima Hi-C+ Kit | Arima | A410321 | Component Box A |
|  |  | A410322 | Component Box B |
|  |  | A410239 | Component Box C |
| Kapa Hyper Prep Kit | Roche | KK8504 |  |
| Ampure XP Beads | Beckman Coulter | A63880 |  |
| Qubit HS DNA Assay | Invitrogen (Thermo) | Q32854 |  |
| Covaris<br>microTUBE | Covaris | 520045 | Full name: Covaris microTUBE AFA Fiber Pre-Slit Snap-Cap<br>6x16mm |
| Illumina TruSeq<br>Adapters | Illumina | 20015960 | Full name: Illumina TruSeq DNA Single Indexes Set A; NOW<br>DEFUNCT |
| Illumina TruSeq<br>UDI v2 | Illumina | 20040870 | Replacement for above; used in only most recent Hi-C Lib<br>(E1AF) |
| Kapa Library<br>Quant Kit | Roche | KK4824 | Full name: KAPA Library Quantification Kit (Illumina)<br>Lightcycler 480qPCR Mix |

### Hi-C sequencing data processing

Sequencing data were obtained in the FASTQ format. These data were then analyzed using the Juicer pipeline. mRatBN7.2/rn7 was used as the reference genome in this analysis. The analysis workflow is provided in our github repository.

## Data Records

The Hi-C sequencing data comprise three groups: (1) four individual rats representing the parental strains of the two recombinant inbred (RI) strains of the hybrid rat diversity panel 2) five RI strains and (3) one F1 hybrid offspring (SHR/Olalpcv x BN/NHsdMcwi). These datasets, containing paired-end raw sequencing reads in fastq.gz format, have been deposited in the Sequence Read Archive (SRA) under accession numbers shown in Table 2.

**Table 2.**
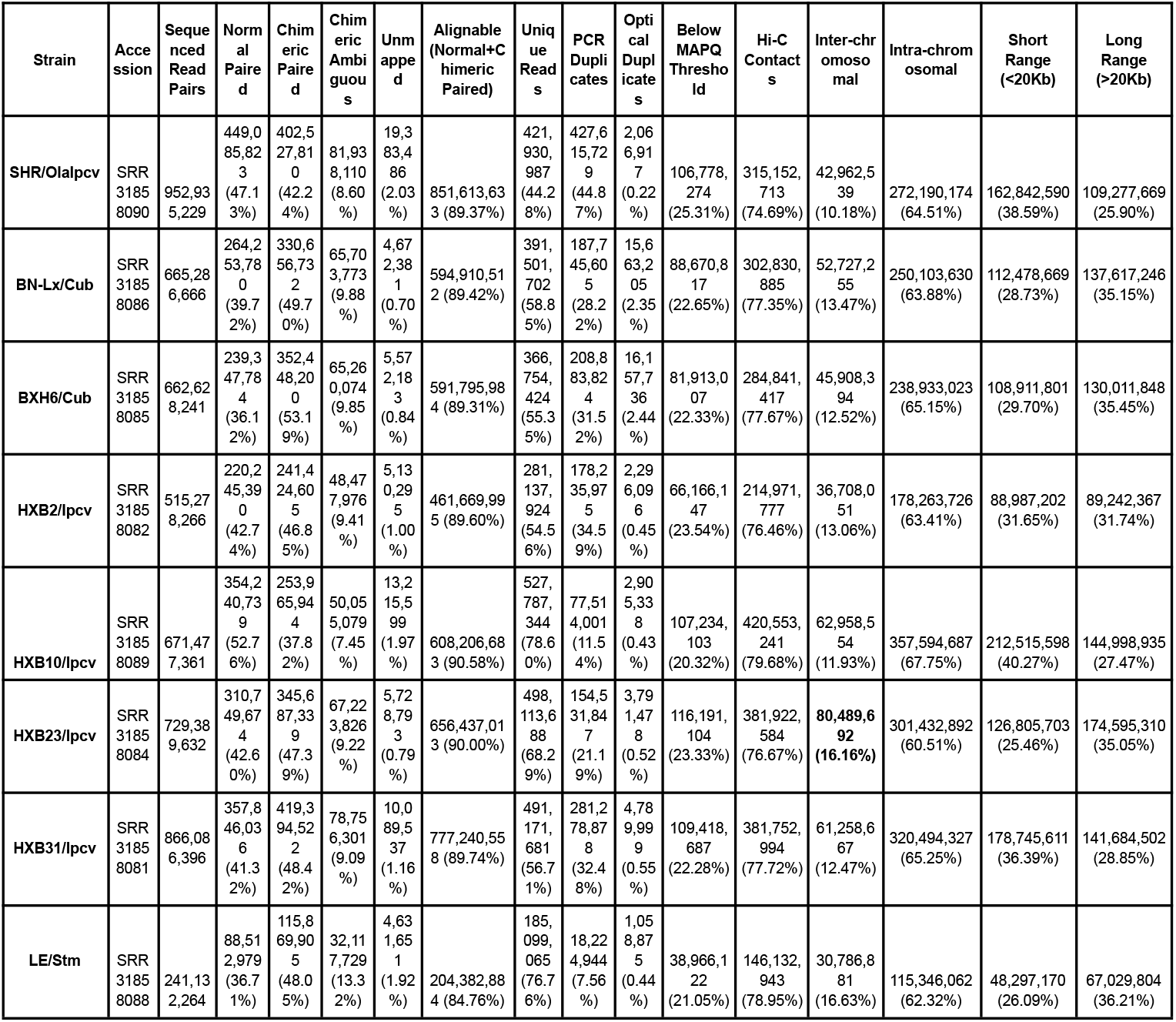
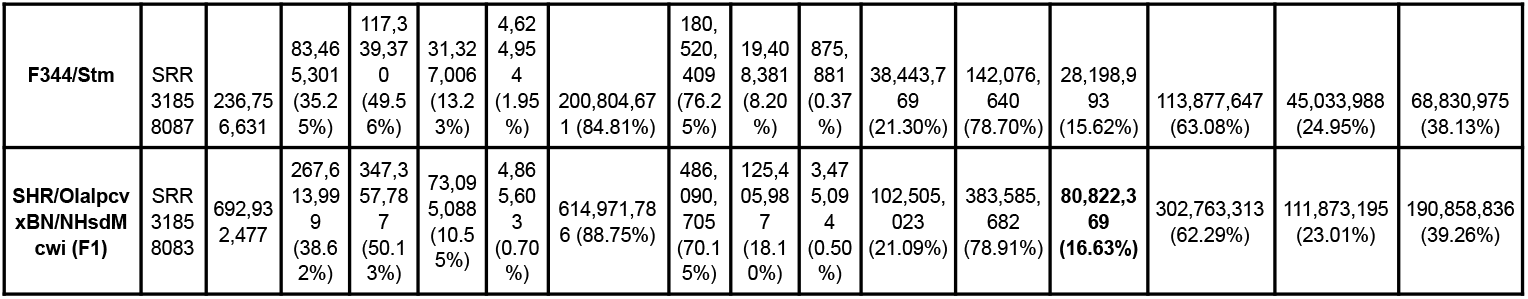
Statistics of Hi-C sequencing after running the Juicer pipeline.

## Technical Validation

We analyzed the Hi-C data using the Juicer pipeline^12^. The mRatBN7.2/rn7 was used as the reference genome. The QC values for each sample were provided in Table 2. We aimed to obtain about 500-600 M paired-end reads per sample. The average reads per sample was 623.39 M reads in our data collection. Two samples, LE/Stm and F344/Stm, fell short of this goal. On average, only 1.31% was not mappable to the reference genome. Among the mapped reads, 41.30% were normal reads that did not contain segments from two different genomic locations (i.e., chimeric reads), 47.34% were chimeric and were mapped to a unique location in the reference genome. The remaining 10.06% were chimeric and matched to multiple locations in the reference genome. For the alignable reads (i.e., normal+chimeric paired), 0.83% were optical duplicates, and 23.83% were PCR duplications, the remaining 63.98% were unique reads. On average, 22.32% of the unique reads mapped to the reference genome with a low quality (MAPQ score less than 30), the remaining 77.68% (297.38 M reads) had sufficient map quality and were used to generate information on Hi-C contacts. Among these reads, on average, 52.28 M were mapped to between chromosomal interactions, and the remaining 245.10 M reads per sample were mapped to within chromosomal interactions. These within chromosomal reads were further divided by the distance between the two interacting chromosomal locations. About 119.65 M reads per sample were from short range (<20kB) interactions, and the remaining 125.41 M reads were from long range (>20 Kb) interactions. Together, these analyses indicate that all our samples pass the quality recommendations put forth by Arima Genomics. These metrics include less than 6% unmapped reads, less than 20% chimeric ambiguous reads, greater than 80% alignable reads and approximately 20% interchromosomal reads. We anticipate this high quality dataset will contribute to the understanding of 3D genomic structure and functional genomics of the rats.

## Code Availability

Data processing was conducted using Juicer 1.6 (https://github.com/aidenlab/juicer/tree/main). The command line used and the input files are available on GitHub at https://github.com/distilledchild/rat-cortex-hic-descriptor/blob/main/README.md

## Author Contributions

PK performed data analysis and drafted the manuscript. RRW generated the Hi-C data and contributed to manuscript preparation. HC oversaw data generation and analysis. HC, RWW, and BMS designed the experiments and revised the manuscript.

## Acknowledgments

The authors thank Dr. Melinda R. Dwinell (Medical College of Wisconsin) for providing the breeders for the Hybrid Rat Diversity Panel (HRDP) and acknowledge the Center for Integrative and Translational Genomics at the University of Tennessee Health Science Center for its support in maintaining the HRDP. The authors also acknowledge Mr. Angel Garcia Martinez and Ms. Caroline Jones for their assistance with HRDP breeding. The Molecular Resource Center and the Genomics Core at the University of Tennessee Health Science Center are acknowledged for their contributions to sequencing data generation. The majority of the computation for this work was performed on the University of Tennessee Infrastructure for Scientific Applications and Advanced Computing (ISAAC) computational resources. This work was supported by NIH/NIDA grants U01 DA-053672 (to BMS, RWW, and HC).

